# Influence of Gaze, Vision, and Memory on Hand Kinematics in a Placement Task

**DOI:** 10.1101/2023.09.21.558829

**Authors:** Gaelle N. Luabeya, Xiaogang Yan, Erez Freud, J. Douglas Crawford

## Abstract

People usually reach for objects to *place* them in some position and orientation, but the placement component of this sequence is often ignored. For example, reaches are influenced by gaze position, visual feedback, and memory delays, but their influence on object placement is unclear. Here, we tested these factors in a task where participants placed and oriented a trapezoidal block against 2D visual templates displayed on a frontally located computer screen. In Experiment 1, participants matched the block to three possible orientations: 0° (horizontal), +45° and −45°, with gaze fixated 10° to the left/right. The hand and template either remained illuminated (closed-loop), or visual feedback was removed (open-loop). In Experiment 2, a memory delay was added, and participants sometimes performed saccades (centripetal, centrifugal, or opposite side). In Experiment 1, the hand consistently overshot the template relative to gaze (similar to reaching), especially in the open-loop task. After a memory delay, location errors were influenced by both template orientation and gaze position. Based on previous reach experiments, we expected these errors to be independent of the previous eye position, but placement overshoot also depended on previous saccade metrics. Hand orientation over-rotated relative to template orientation (all conditions). Orientation was influenced by gaze position in Experiment 1, but this vanished in the presence of a memory delay. These results demonstrate interactions between gaze, location, and orientation signals in the planning and execution of hand placement and suggest different neural mechanisms for closed-loop, open-loop, and memory delay placement.

**NEW & NOTEWORTHY:** Eye-hand coordination studies usually focus on object acquisition, but placement is equally important. Here, we investigated how gaze position influences object placement toward a 2D template, with different levels of visual feedback. Like reach, placement overestimated goal location relative to gaze, but was also influenced by previous saccade metrics. Gaze also modulated hand orientation, which generally overestimated template orientation. Gaze influence was feedback-dependent, with location errors increasing but orientation errors decreasing after a memory delay.

## INTRODUCTION

Goal-directed hand movements facilitate a productive interaction with the world around us (1). In the case of prehension (reach to grasp movements), the influence of factors such as gaze position and levels of visual feedback is very well studied (e.g., 2–4). However, the interaction with surrounding objects often does not end at object acquisition (5–7). Take, for example, toasting a flat piece of bread. After acquiring the object, it must be transported, oriented, and correctly placed into a toaster slot. Despite their real-world importance, the determinant of placement performance has been relatively neglected compared to prehension. Here, we examined the influence of gaze position, saccades, and levels of visual feedback (open-loop, closed-loop, memory-guided) on hand location and orientation (and their interactions) in a simple visual / memory-guided placement task.

Grasp and placement share several common characteristics, e.g., they both need to locate and orient the hand, and both are influenced by task complexity (8–10). However, they differ in how they are influenced by sensory information and in their movement intention. First, grasping movements are generally aimed toward a visually perceived object, whereas placement movements could rely on the imaginary image of the movement outcome or involve matching a target template (such as the toast example above). Grasp ends with haptic feedback physical object, whereas placement follows the opposite sequence. Several studies examined sequential coordination with reach and placement, for example, during eating (11–13), and the way they influence each other (14). For instance, the final movement intentions of placement influence movement kinematics during the grasping stage (15–17). However, several fundamental aspects of eye-hand coordination that are well-studied in the reach system have not been studied in placement.

First, it is known that gaze direction strongly influences reaching and pointing. Gazing directly at the target increases our movement accuracy (18, 19). In some cases, gaze remains locked onto the goal during reaching (20), but in other real-world conditions gaze must remain focused on other tasks or anticipates future goals during the current reach (5, 7, 21). When the gaze deviates from the target, people tend to make pointing / reaching errors that ‘overshoot’ in the opposite direction (3, 18, 19). In the reach system, this overshoot is entirely determined by the final gaze position at the time of pointing (independent of previous saccades) consistent with the use of a gaze-centered updating mechanism for the reach goal (2, 3, 22). This is clinically relevant, because these gaze-dependent errors become more pronouced in the presence of parietal cortex damage (23). It is noteworthy that these topics have mainly been studied using centrifugal saccades away from the target (2, 3, 24), so it is not clear the results extend to peripheral-to-peripheral or centripetal saccades (25). To our knowledge, these basic eye-hand coordination issues have not been investigated in a placement task.

Second, a number of grasp studies have examined the kinematics of orienting the hand to the object (26, 27). Grasp orientation involves the coordination of the entire hand-arm system (28) but can be dissociated from reach location at both the behavioral and neural levels (29–31). Place-to-match tasks show an oblique effect where oblique targets show an increase in orientation error in comparison to their cardinal counterpart (32, 33). In contrast to pantomiming movements, placement orientation is immune from damage to the ventral visual stream, suggesting a dorsal (parietal) mechanism (34). In the presence of saccades, grasp orientation is also dependent on final gaze position (27), and this is reflected in gaze-grasp orientation interactions at the level of the parietal cortex (35). Again, none of these factors have been studied in placement movements, so it is important to show how gaze and hand orientation interact in healthy participants, not only to understand the system, but to establish a normal baseline for patient studies.

Importantly, the preceding descriptions of behavioral errors are highly dependent on levels of visual feedback and memory. In the absence of allocentric cues (such as visual landmarks in the surround), movement errors generally increase as the delay between target presentation and movement onset increases (36–38). Visual feedback is particularly important for the final phase of action (39), although orientation errors can also be improved by proprioceptive information about the target (32). Likewise, in the case of the gaze-dependent reach and orientation errors described above, these are normally overridden by visual feedback (2, 3, 27). There are exceptions to these rules: for example, illusion-dependent grasp errors tend to dissipate with a memory delay (34, 40). And the larger gaze-dependent errors seen in optic ataxia may dissipate over time, suggesting different spatial mechanisms for immediate versus delayed reaching. The overall point is that reach and grasp errors may be highly sensitive to different levels of visual feedback, ranging from full visual feedback of the hand and desired goal (closed-loop) to the removal of this feedback (open-loop) to the addition of a memory delay.

The purpose of the current study was to investigate how hand location and orientation are influenced by gaze position, previous saccades, and various levels of visual feedback (closed-loop, open-loop, memory delay) in a placement task. Our hypothesis was that if the placement system shared similar mechanisms as the reach system (in the absence of proprioceptive feedback), we should see gaze-dependent overshoots errors in placement location that should be independent of previous saccades, and likewise, any gaze-dependent hand orientation errors should only depend on final gaze position. We further expected these errors to grow with the progression from closed-loop to open-loop to memory delay, as the system has to increasingly rely more on noisy internal models without the opportunity for visual feedback corrections.

To test this, we asked participants to place a trapezoidal shape against an oriented template in the centre of a computer screen, with their gaze fixated on the template or deviated to the left or right. We divided this into two experiments, where 1) closed-loop and open-loop performances were compared, and 2) a memory delay was added to the open-loop condition, accompanied by sustained fixation or a saccade (which could be centripetal, centrifugal, or across to the template). We also tested interactions between hand location and orientation on these tasks. The results showed that our task yielded some of the predicted effects (such as an open-loop gaze-centered overshoot), but other results went contrary to expectation, such as reduced orientation errors after a delay.

## METHODS

### Participants

A total of 36 young adults participated in this study, with 17 individuals taking part in Experiment 1 (8 females, 9 males, 14 right-handed, ages 22 - 34) and 19 in Experiment 2 (7 females, 12 males, all right-handed, ages 19 - 28). Notably, 5 participants completed both two experiments. Of these, 16 (Experiment 1) and 14 (Experiment 2) participants, respectively, were included in our data analysis (see Data Analysis and Exclusion Criteria) providing sufficient power for our analysis (see Statistical Analysis). Each participant exhibited normal or corrected-to-normal visual acuity, with no reported neurological disorders. Written consent was collected from all participants, and they received monetary compensation for their time. All experimental procedures were approved by the York University Human Participants Review Committee.

### Apparatus and Stimuli

A 70 cm × 40 cm Sony Television screen was used to present visual stimuli, which included gaze fixation points (a white dot to the right, left or centre of the screen), and rectangular target templates (a white rectangle display at the centre of the screen in three possible orientations). This screen was mounted onto a stable table and connected to a computer for stimulus control (Fig. 1). The screen was covered with dark tints to decrease luminance (luminance = 0.000 cd/m^2^ measured with Konica Minolta® Luminance Meters LS-100) and minimize extraneous allocentric cues. A chinrest, placed at the edge of the table, maintained the participant’s head positioned 26 cm away from the screen and served to reduce body movements. The participant sat on an adjustable chair with their head on the chinrest and was positioned with their right eye directly in front of the centre of the screen. A green LED light depicted the home position for the hand, halfway between their torso and the screen on the platform.

**Figure 1:**
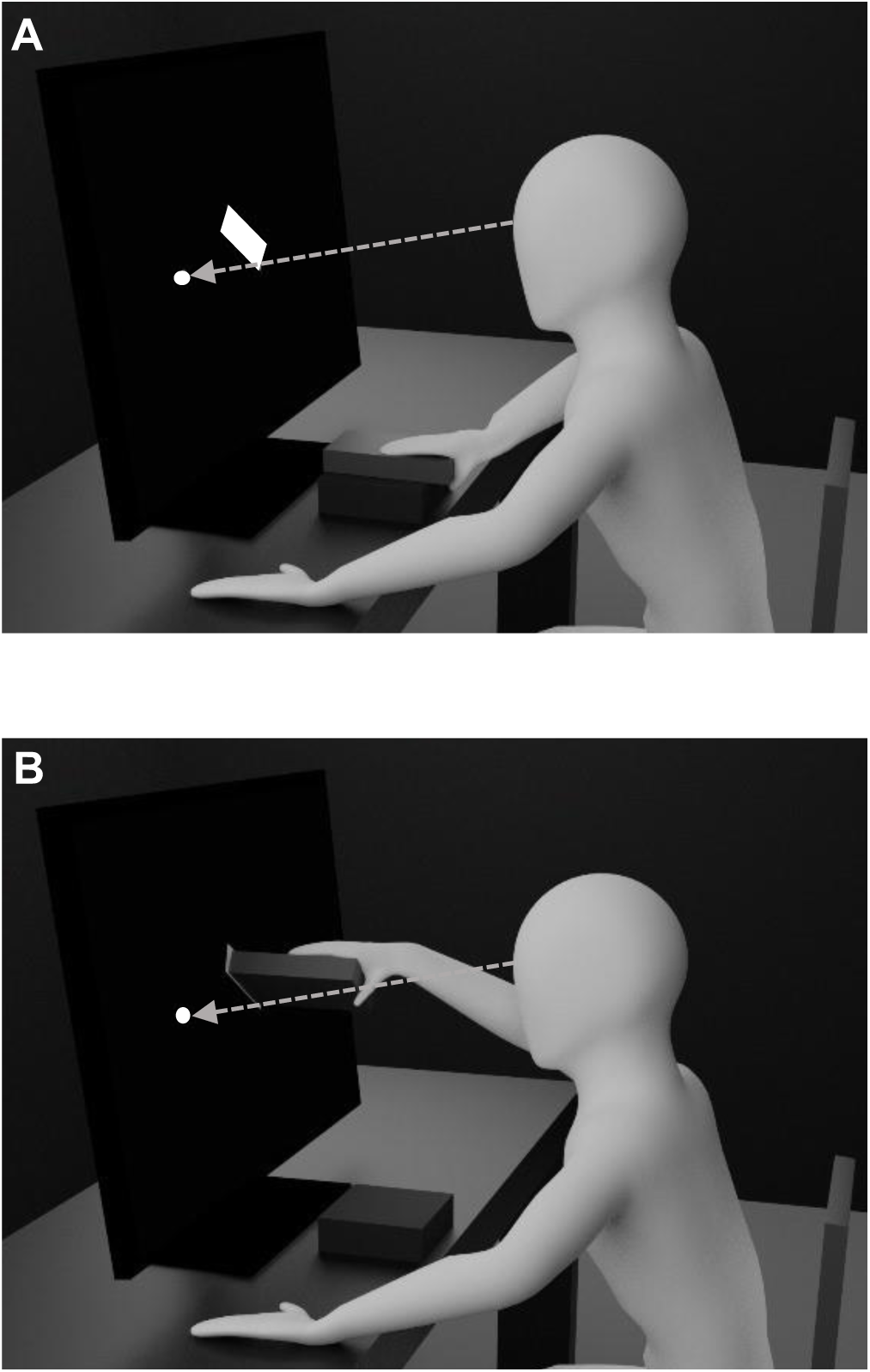
Experimental Task. A) Depicts the preparatory period before the go signal, with the hand at the resting position, gaze on the fixation (white dot to the left) and the target template (white rectangle) displayed at the screen center. B) Example of a final object placement against the template, while gaze is maintained at the fixation point. The chinrest and eye-tracking device are not shown in this illustration.

The task involved placing a trapezoidal block against a template displayed on the computer screen (see next section for details). In its horizontal position, the trapezoidal block had a height of 2.0 cm, a width of 7.5 cm, with a depth of 10.0 cm on the left side, and 8.0 cm on the right side. An elastic band passed through two openings and was tied around the participant’s fingers. The object’s real-time dynamic and kinetic motions were tracked and analyzed using the Optotrak 3020 motion capture system (Northern Digital Inc, Waterloo, On, Canada) at a sampling rate of 100 Hz; the system provided measurements of the hand position and motion during the experiment. The system monitored the position and motion of the block by collecting the location of six InfraRed Light Emitting Diodes (IREDs), which were affixed in groups of three onto two surfaces of the prism (the top surface and the left adjacent surface). They were used to create two three-dimensional virtual points that were average to calculate the central position of the front surface of the object. The gaze location of the left eye was monitored using ISCAN Eye Tracking system (ISCAN Inc, Woburn, Massachusetts, United States) which was equipped with a camera and an infrared light source. The camera captured the infrared reflection of pupil and conea, based on which, the system software calculated the 2D gaze after the calibration. The analog outputs of the gaze signals were sent to Optotrak Data Acquisition Units (ODAU). With a sampling rate of 1000Hz, the digitized 2D gaze data was recorded together with Optotrak IREDs data during the experiment.

### General Task Design

We conducted two psychophysical experiments to gain a fundamental understanding of how object placement is influenced by gaze location, target orientation, and level of visual feedback. The study investigated how varying levels of visual feedback (in Experiment 1) and the presence of saccades (in Experiment 2) affected the endpoint positioning of an object to match the location and orientation of the target. In the two tasks, the design included three gaze location / fixation points either at the center of the screen, to the left, or to the right of the central location. The target, a rectangular object template, was always presented at the center of the screen and could have either horizontal (0°), +45°, or −45° orientation (Fig. 2A). These orientations were chosen based on the range that participants found comfortable in a preliminary testing session.

**Figure 2:**
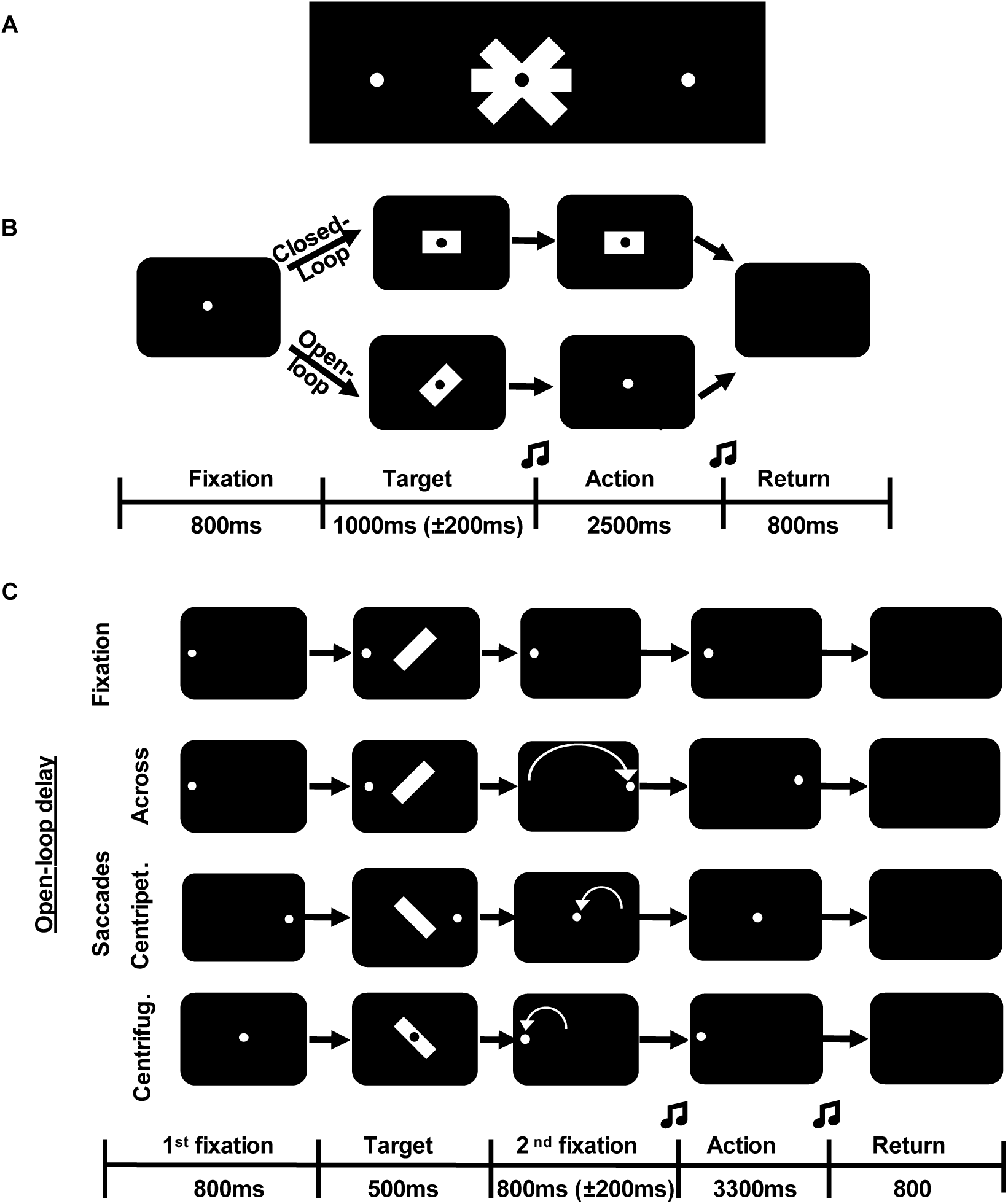
Experimental protocol. A) Illustrates the 3 possible fixation locations (left, center, and right) and target template orientation (−45°, 0°, and +45°) presented on a black LED screen; B) Schematic sequences of events for *Experiment 1*. Black rectangles represent the stimulus screen at different time points in a trial. Initially, the screen was black. Then, the gaze fixation stimulus appeared (in this case at the center). In the *Closed-Loop Condition*, the target template (horizontal in this example) remained illuminated until an auditory ‘go signal’ indicating action onset. In the *Open-Loop Condition*, the target template (+45° in this example) disappeared as the signal to initiate action. In both cases, the participant placed and maintained the object against the object template until an auditory signal indicated return to the home position. C) Schematic sequence of events for memory and saccade conditions in *Experiment 2*. Participants first aimed gaze toward the initial fixation point (left, right and centre depending on the condition). Then, a +45° or −45° target template is presented for 500ms, followed by an 800ms memory delay. Depending on the task, participants then either maintained fixation during the delay (*Fixation Condition*), made saccades across to the opposite fixation stimulus (*Saccade across*), saccades away from centre fixation (*Centrifugal*), or toward the central fixation (*Centripetal*). Afterwards, an auditory ‘go signal’ instructed participants to place the object to the remembered target template orientation.

At the beginning of each trial, all room lights were extinguished, and participants placed the trapezoidal block at the home position in a horizontal orientation with the right hand (fingers above, thumb below). Participants then fixated their gaze on a point that appeared at one of the three positions (Fig. 2A). The rectangular target template (slightly larger than the block’s forward 2 × 7.5 cm surface) then appeared at the centre of the screen (aligned with the centre fixation point). Participants were instructed to place the block orthogonal to the screen (with various levels of visual feedback) such that the forward pointing section aligned with the location and orientation of the template (Fig. 1). Finally, they returned the hand / block to the home position and prepared for the next trial, (the detailed sequences of stimulus events are provided in the following two sections.)

### Experiment 1

Experiment 1 investigated the influence of gaze location and visual feedback on integrating a stimulus location and orientation during an object-placement task. The experiment was a within-subjects design involving 3×3×2 study parameters (three gaze positions, three orientations, and two visual conditions). The two visual feedback levels were either closed-loop with full visual feedback of the target template and the hand or open-loop with no visual feedback of the target template and the hand.

In all trial sequences (Fig. 2B), participants were allowed 800ms with nothing on the screen to hand to settle their hand position and prepare for the trial. Then, one of three fixation dots (centre, 13° visual angle left or right) appeared and remained on for the entire duration of the trial. Participants maintained fixation on this dot throughout the trial. 800ms after the initial presentation of the fixation dot, a rectangular target template appeared at one of the three possible orientations. This target template was displayed for 1000±200ms before a beeping sound cued the placement movement. In the closed-loop condition, the target template remained visible, and light illuminated the hand and object for the duration of the movement; whereas in the open-loop condition, the target template was extinguished in an otherwise dark room (except for the fixation point). In both cases, participants had 2500ms to execute the movement and maintain the object placed against the screen until a return beeping signal was played. Then, the home LED was illuminated, and they had 800ms to return to the home position. This was followed by a 2100ms intertrial interval when two light bulbs behind the monitor were illuminated to prevent dark adaptation.

In each session, participants completed a total of 390 trials divided into five runs of 78 trials. Before starting a run, participants completed an eye calibration. Each run began and ended with three calibrations (closed-loop) trials, which were only used offline to check the stability of the recording signals. The order of conditions was pseudorandomized so that each of the 18 possible trial types (three orientations × three gaze positions × two feedback conditions) was repeated 20 times across the entire experiment. Each run was 10.4 minutes long. Participants were given two minutes to rest after each run. Each experiment lasted approximately 72 minutes.

### Experiment 2

The 2^nd^ experiment investigated the influence of saccade on placement location and orientation. The experiment was a within-subjects design involving 3 (Initial Gaze Locations) × 3 (Final Gaze Locations) × 2 (Orientations) study parameters. Here, there was always a memory delay (with no visual feedback of the target template or the hand) between seeing the target template and placing the object. The task design included three gaze locations (center screen, 10° visual angle left or right). Here, the fixation point could change (2/3 of the time) after the target template presentation, causing a saccade to occur after the target template disappeared and the change in fixation location prone participants to do a saccadic eye movement. Combining the three gaze locations as initial and final fixation positions gave six types of saccadic eye movements (centrifugal left or right from centre, centripetal from left or right, or across from left to right and vice versa which involved a shift from one visual field to the other). The target template was presented in two possible orientations (+45° or −45°). Participants were presented with the target template and performed the placement movement after a brief memory delay.

Experiment 2 followed a similar sequence as Experiment 1 (Fig. 2C). The initial fixation point in the three possible locations (left, centre, or right) appeared 500ms after the trial started. After 800ms, the rectangular target template was presented for 500ms. As the target template was extinguished, the fixation point either remained in the same location (servings as open-loop delay) or moved to a new horizontal location for the saccade conditions. A beeping sound was played 800±200ms later and served as a cue to initiate the placement movement toward the TV monitor. Participants had 2500ms to perform the movement and maintained the object placed against the screen until a return beeping signal was played. Then, they had 800ms to return to their home position, which was illuminated during that period. The return time is followed by a 1000ms intertrial interval when two light bulbs behind the monitor are illuminated to prevent adaptation to the darkness.

In each session, participants completed a total of 432 trials divided among six runs of 72 trials. At the start of each run, participants completed an eye calibration. The order of conditions was pseudorandomized so that the 18 possible trial types (three initial gaze positions × three final gaze positions × two orientations) were repeated 24 times across the entire experiment. Each run was 8.28 minutes long. Participants were given two minutes to rest after each run. Each experiment lasted approximately 72 minutes.

### Data Analysis and Exclusion Criteria

Eye-tracking and hand-tracking data were exported from the Optotrak software and presented into a custom graphical user interface written in MATLAB (MathWorks Inc., Natick, MA). The data were preprocessed to ensure subjects’ compliance with the procedure. The program automatically marked the onset of movement from the home position and recorded the movement endpoint when the object is placed in the target template location. The data was recorded in millimeter coordinates. Each trial went through a series of four exclusion criteria. The first exclusion criterion required participants to maintain fixation to the target template gaze location. The second exclusion required that they reached their target template alignment before the return signal was played and the fixation point disappeared. The third criterion required that trials be discarded when the participant’s movement onset occurred before the beep signal in memory-guided trials. The fourth criterion was enforced when the participant’s orientation deviated by more than ±25° from the target templates. The inability to satisfy either one of these criteria resulted in the omission of the trial for further analysis. In Experiment 1, one participant’s data was excluded because they failed to maintain fixation while performing hand movements in multiple trials. In Experiment 2, data from five participants’ data were excluded because they did not attain the minimum number of valid trials for 1/3 of the conditions. For participants included in the analysis, 76.2% of the trials were used for Experiment 1 and 70.4% for Experiment 2.

### Statistical Analysis

For each valid trial, the endpoint orientation and horizontal position were compared to the actual target template orientation and horizontal position (Fig. 3 *Top Row*). Statistical analyses of each condition for each participant were conducted on the mean error to obtain placement movement accuracy in endpoint horizontal location and in endpoint orientation. In Experiment 1, to investigate how gaze location, target template orientation, and visual guidance affected movement accuracy, a three-way repeated-measures analysis of variance (RM ANOVA on three orientations (0°, −45°, and 45°) × three gaze locations (right, centre and left) × two visual guidance (visually or memory-guided actions)) was conducted on accuracy the endpoint orientation and the endpoint horizontal location. In Experiment 2, to investigate how adding a memory delay and different types of saccades influenced movement accuracy, a three-way repeated-measures analysis of variance (RM ANOVA on three presaccade gaze positions (right, centre and left) × three postsaccade gaze positions (right, centre and left) × two target template orientations (−45° and 45°) was conducted on accuracy the endpoint orientation and the endpoint horizontal position. All statistical analyses were conducted in RStudio software. A post-hoc power analysis was also conducted to confirm the adequacy of our sample size was large enough to capture the effect for a two-tailed test with power at 0.8 and alpha of 0.05. We found that a sample size of 10 was determined to be sufficient for detecting the main effects. Mauchly’s test was used to assess if sphericity was violated. In cases where sphericity was not assumed, Greenhouse-Geisser’s correction was used. If the RM ANOVA was significant with a p < 0.05, we conducted post-hoc tests to adjust for multiple comparisons. A follow-up interaction contrast test was conducted in cases where interaction was reported.

**Figure 3.**
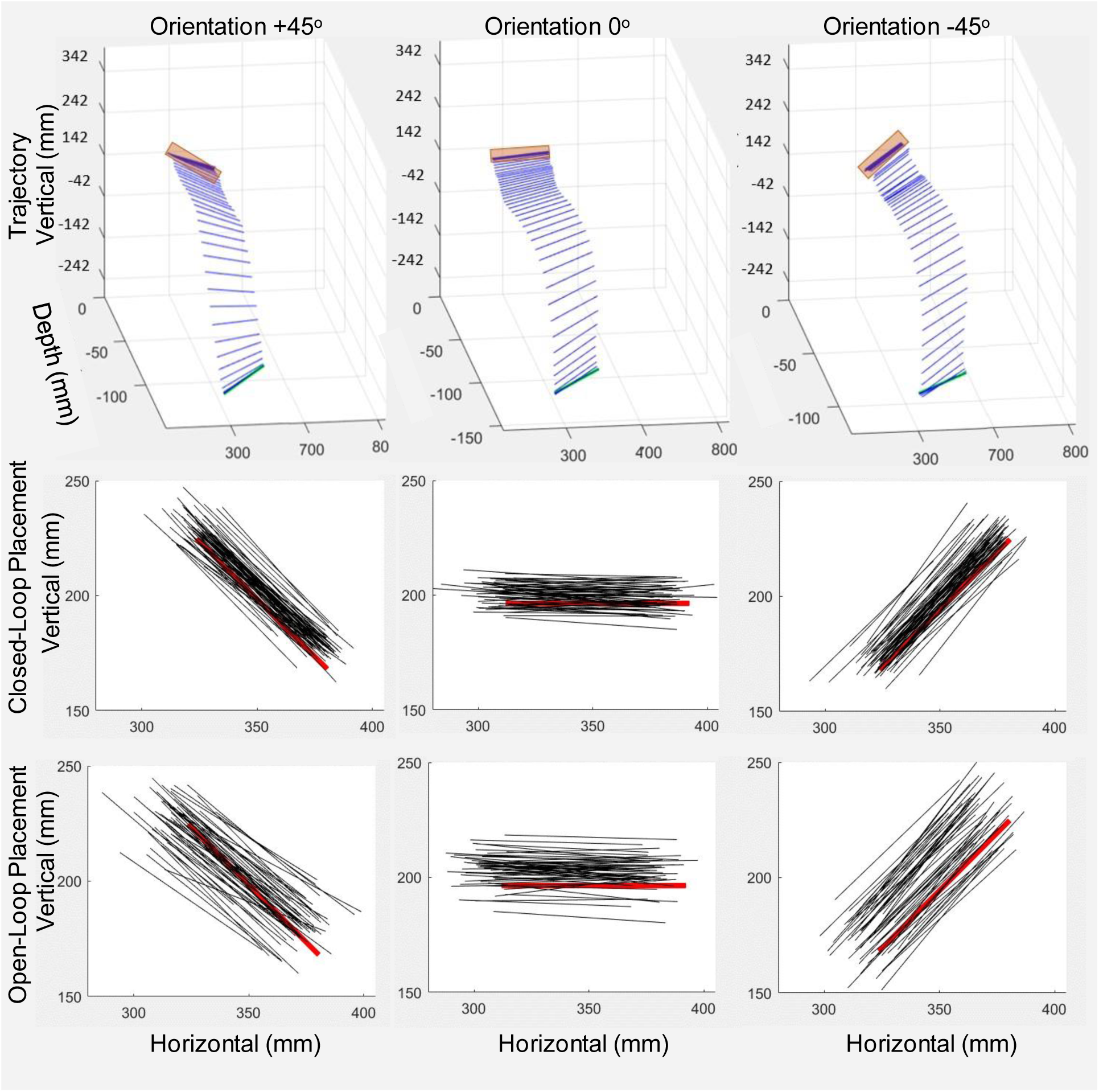
Typical placement kinematics for one participant in Experiment 1. The *top row* shows example trajectories of the forward-facing side of the object (black lines) towards three possible template orientations (pink rectangles, from left to right: −45°, 0°, +45°) in 3D space. The *middle row* shows 2D endpoint orientation and location of object placement (black lines) for all trials for this participant, relative to the template location and orientation (red lines) in the closed-loop condition. The *bottom row* shows the same for the open-loop condition.

## RESULTS

### Experiment 1

Experiment 1 tested how object placement is influenced by gaze location, target template orientation, and level of visual feedback. The study investigated how closed-loop and open-loop conditions during a placement movement influenced the endpoint positioning of an object to match the location and orientation of the central target template. Two sets of data from the same movement were studied: endpoint error in location (relative to the centre point of the target template) and the endpoint error in orientation (relative to the orientation of the template). These two values were defined by comparing the discrepancy between final object position and orientation (after placement) relative to the target template’s actual position and orientation (Fig. 3 *Middle and Bottom Rows*).

### Location error

Error in final object position were measured and analyzed to study the influence visual feedback, gaze position, and target template orientation on horizontal object placement position. The violin plots in Fig. 4 A and *B* summarize the distribution of placement location errors in each condition. Because the physical variation in gaze was in the horizontal plane, we exclusively considered the error in the endpoint position in the horizontal dimension, where a negative value indicates movement to the left and a positive value indicates movement to the right.

**Figure 4.**
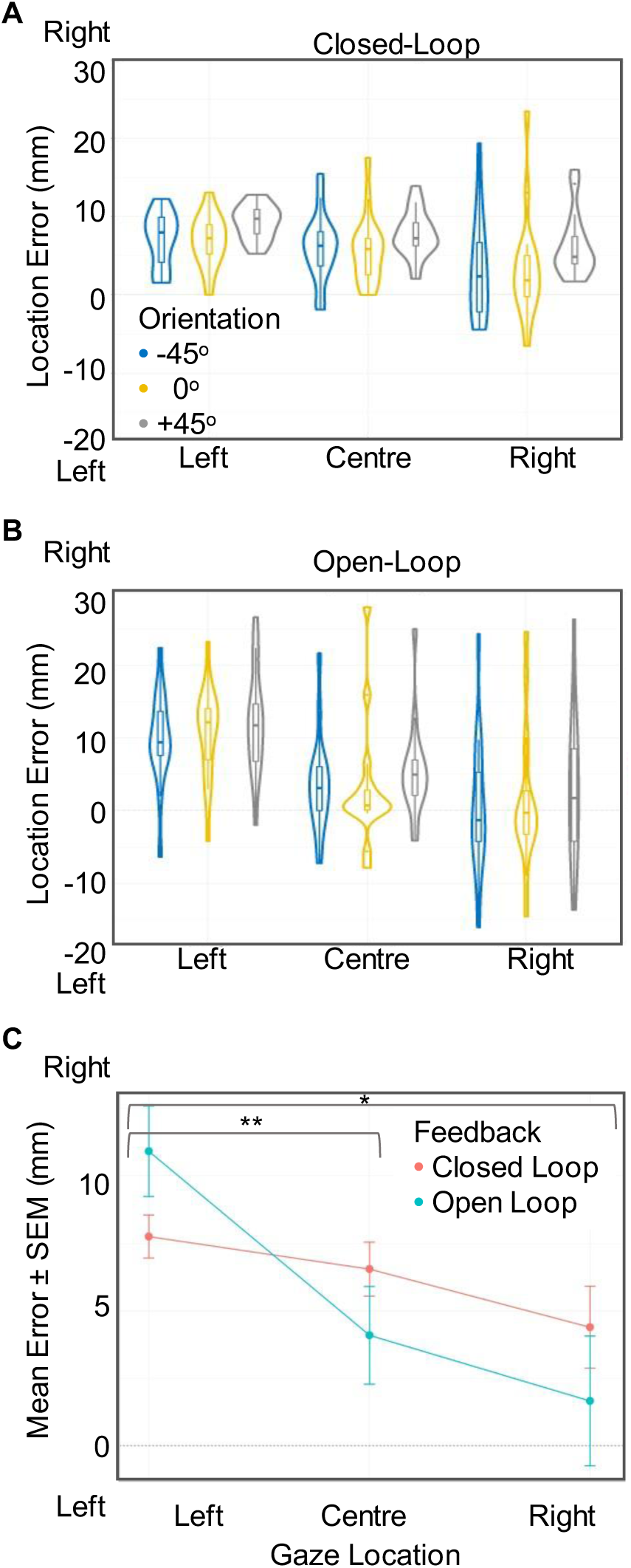
Graph of Location Errors in *Experiment 1*. A-B) Violin plots representing placement location errors for the left, centre, and right gaze fixation locations in the *Closed-Loop* (A) and *Open-Loop* (B) conditions. Violin plots show the distribution of the participant’s average location error and include the median (as the dot) and the interquartile range (as the bar in the center of the violin). Location error was quantified as the horizontal difference between the object centre and template centre, where the mean of participants’ endpoint position is compared to the target template position, and the error is used to compute the accuracy in the horizontal position. The values from the y-axis provide information regarding the direction of the error: a positive value indicates movement errors toward the right, and a negative value indicates movement errors to the left. C) Average Mean horizontal endpoint error (± SEM) across participants in endpoint horizontal location plotted as a function of gaze position. The graph displays the mean error and standard error of participants’ endpoint position on the screen in comparison to the actual target template position on the screen. The x-axis represents the three possible gaze locations (left, centre or right). Asterisks indicate significant gaze-dependent errors, and there was a statistically significant 2-way interaction between gaze location and level of visual feedback.

The data (Fig. 4, A and B) revealed a general error bias to the right, i.e., when looking straight ahead and placing the object to the central target template. There was also a greater spread in location error in the open-loop conditions in comparison to closed-loop guided movements. When comparing the average position error across participants, we could observe a tendency to overshoot location relative to gaze, i.e., they made placement errors to the left when looking to the right and errors to the right when looking to the left (Fig. 4C). In addition, this pattern became stronger in the absence of visual feedback.

We conducted a three-way repeated-measures analysis of variance (ANOVA) composed of 3 (Gaze Locations) × 3 (Orientations) × 2 (Levels of Feedback). The results showed main effects of Gaze Location (F_2,30_ = 8.07, P_GG_ < 0.01, η^2^ = 0.35) and Orientation (F_2,30_ = 7.48, P < 0.01, η^2^ = 0.33) on object location, as well as a significant interaction effect of Location and Levels of Feedback (F_2,30_ = 7.59, P_GG_ < 0.01, η^2^ = 0.34). There was no significant interaction effect between gaze location and orientation (F_4,60_ = 1.033, P = 0.39) and between level of visual feedback and the orientation (F_2,30_ = 0.30, P = 0.74) and no significant three-way interaction effect was found for the endpoint location error (F_4,60_ = 0.56, P = 0.68, η^2^ = 0.04). While there was a bias toward placing the object to the right, the main effect of gaze location showed a significantly greater location error toward the right for leftward gaze locations compared to central and right gaze location respectively. In addition, there was a greater error toward the left in the +45° in comparison to the 0° orientation and the −45° orientation.

A post-hoc test investigating the interaction contrasts between Gaze Location and Level of Visual Feedback revealed a difference in the slope pattern of placement location as a function of gaze location. Bonferroni’s adjustment was employed to control for family-wise error. Because of the overall shift of these placement location errors to the right, the increased gaze-dependent effect (with no visual feedback) caused pointing errors to increase at left gaze position (mean difference = 3.16mm, d = 0.60) but decrease at center (mean difference = −2.46mm, d = 0.42) and right fixations (mean difference = - 2.73mm, d = 0.34).

Overall, participants showed a bias in object placement to the right, and tended to overshoot the central target template relative to gaze position, especially in the absence of visual feedback. Furthermore, the standard deviations showed more variability in the data in the absence of visual feedback. These results tend to agree with observations made in pointing and reaching studies (.e.g., 2, 3).

### Orientation error

Errors in final object orientation were measured and analyzed to study the influence of visual feedback, gaze position, and target template orientation on placement orientation. The violin plot in Fig. 5A summarizes the distribution of orientation errors in each condition, where a negative value indicates a clockwise error in orientation and a positive value indicator counterclockwise error in orientation. With the one exception of the left gaze in the closed-loop condition, participants tended to make more negative (clockwise error) orientation for the −45° template and more positive (counterclockwise) error in the +45° (Fig. 5A). In other words, participants rotated the object too far relative to the target template orientation. Additionally, error distributions tended to be larger in the open-loop condition compared to closed-loop visual feedback.

**Figure 5.**
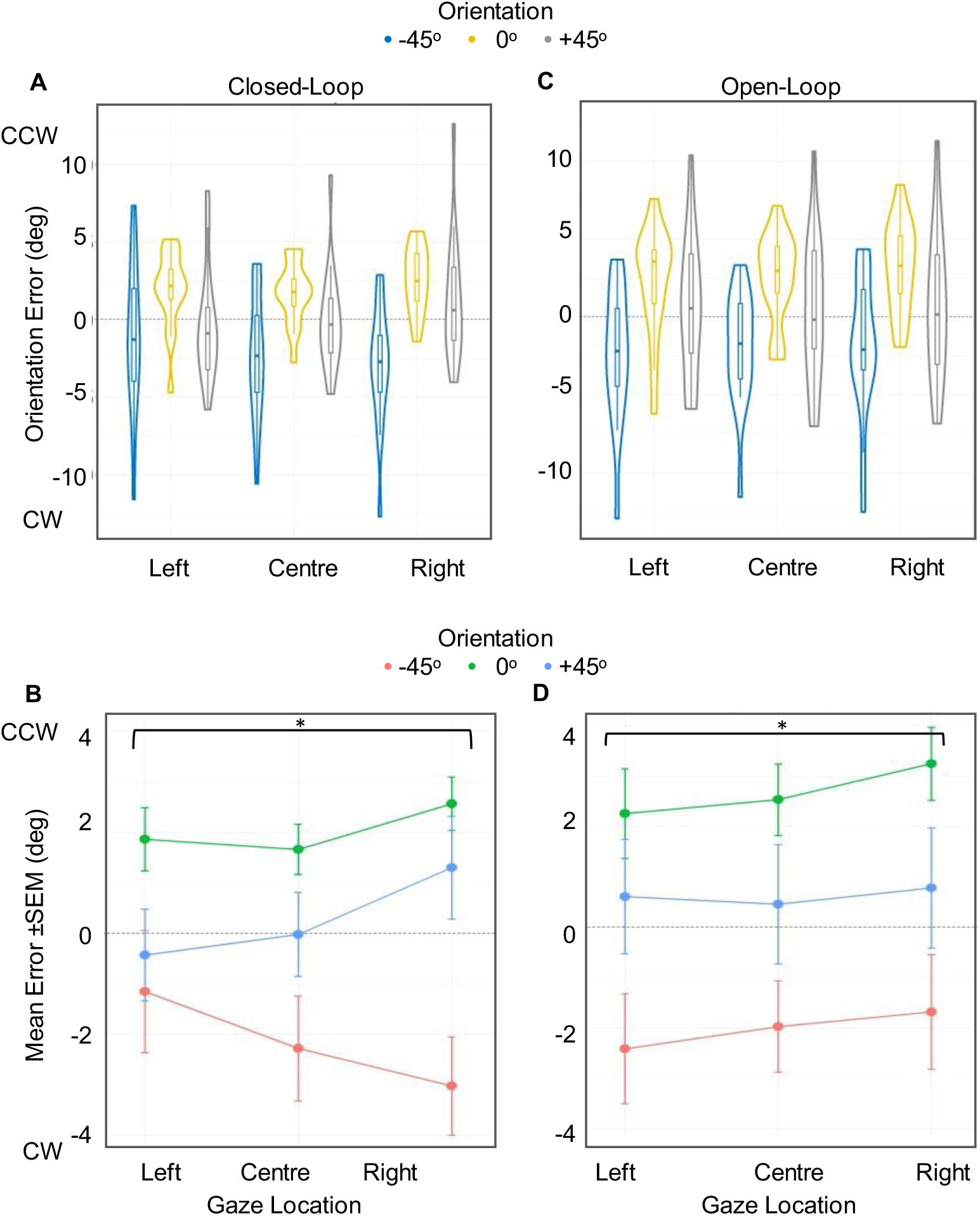
Orientation Errors in *Experiment 1*. A) Violin plots illustrating placement orientation errors for the left, centre, and right locations in the *Closed-Loop* and *Open-Loop* conditions. Violin plots show the distribution of the participant’s average orientation errors and include the median (as the dot) and the interquartile range (as the bar in the center of the violin). Errors were the angular difference between object and template orientation, where a positive value indicates a counterclockwise rotation and a negative value indicates a clockwise rotation. B) Error in endpoint orientation (mean ± SEM across participants) plotted as a function of gaze location. There was a statistically significant 3-way interaction between gaze location, level of visual feedback, and target template orientation. There was a statistically significant 2-way interaction between gaze location and level of visual feedback. Asterisks indicate significant gaze-dependent errors for +45° and −45° such that the orientation errors were in opposite directions depending on the gaze location in the closed-loop condition and the tendency to rotate more clockwise for the left gaze and more counterclockwise for the right gaze in the open-loop condition.

Figure 5. Orientation Errors in *Experiment 1*. A) Violin plots illustrating placement orientation errors for the left, centre, and right locations in the *Closed-Loop* and *Open-Loop* conditions. iolin lots s ow t e distri ution o t e artici ant’s average orientation errors and include the median (as the dot) and the interquartile range (as the bar in the center of the violin). Errors were the angular difference between object and template orientation, where a positive value indicates a counterclockwise rotation and a negative value indicates a clockwise rotation. B) Error in endpoint orientation (mean ± SEM across participants) plotted as a function of gaze location. There was a statistically significant 3-way interaction between gaze location, level of visual feedback, and target template orientation. There was a statistically significant 2-way interaction between gaze location and level of visual feedback. Asterisks indicate significant gaze-dependent errors for +45° and −45° such that the orientation errors were in opposite directions depending on the gaze location in the closed-loop condition and the tendency to rotate more clockwise for the left gaze and more counterclockwise for the right gaze in the open-loop condition.

A second three-way ANOVA was conducted to investigate the influence of gaze location, target template orientation, and visual feedback on final object orientation during placement movements. The ANOVA revealed a significant main effect of Orientation (F_2,30_ = 6.96, P_GG_ < 0.01, η^2^ = 0.32), a significant two-way interaction effect between Orientation and Gaze Location (F_4,60_ = 3.39, P < 0.05, η^2^ = 0.18) and a three-way interaction between Orientation, Gaze Location, and Visual Feedback (F_4,60_ = 9.48, P < 0.0001, η^2^ = 0.39).

Post-hoc tests were conducted to investigate the three-way interaction contrasts between gaze location, target template orientation, and visual feedback levels. After correction for multiple comparisons, there was a significant difference in the slopes when comparing the error in orientation for the +45° and – 45° degree angles in the two levels of visual feedback when the gaze location was to the left or the right (P < 0.05). Therefore, both gaze location and visual feedback levels influence error in orientation for the oblique orientations (Fig. 5B).

In summary, participants tended to over-rotate the block relative to the target template, but this was modulated by gaze position and visual feedback. For example, the right gaze location showed greater rotation error than the left gaze location in the closed-loop condition, and the reverse trend in the open-loop condition.

### Experiment 2

The main purpose of Experiment 2 was to 1) assess object placement location and orientation errors when a memory delay was added to the task, and 2) examine how these errors are influenced by the occurrence of different types of saccades (centripetal, centrifugal, across) during this memory interval. In particular, we asked whether errors were influenced by the final gaze position alone or were also influenced by the initial gaze position / saccade vector. Once again, we analyzed the endpoint accuracy error in location (relative to the centre point of the visual template) and the endpoint accuracy error in orientation (relative to the orientation of the template) (Fig. 3 *Middle Row*).

#### Location error

The violin plots in Fig. 6A/B show the distributions of error in placement relative to the target template for each template orientation and final gaze position. Overall, the errors tend to overshoot the target relative to gaze; for instance, they lean more to the right (positive) for leftward gaze. They are also slightly modulated by template orientation: there is a tendency to place the object more left for clockwise (negative) orientations and more right for counterclockwise (positive) template orientations.

**Figure 6.**
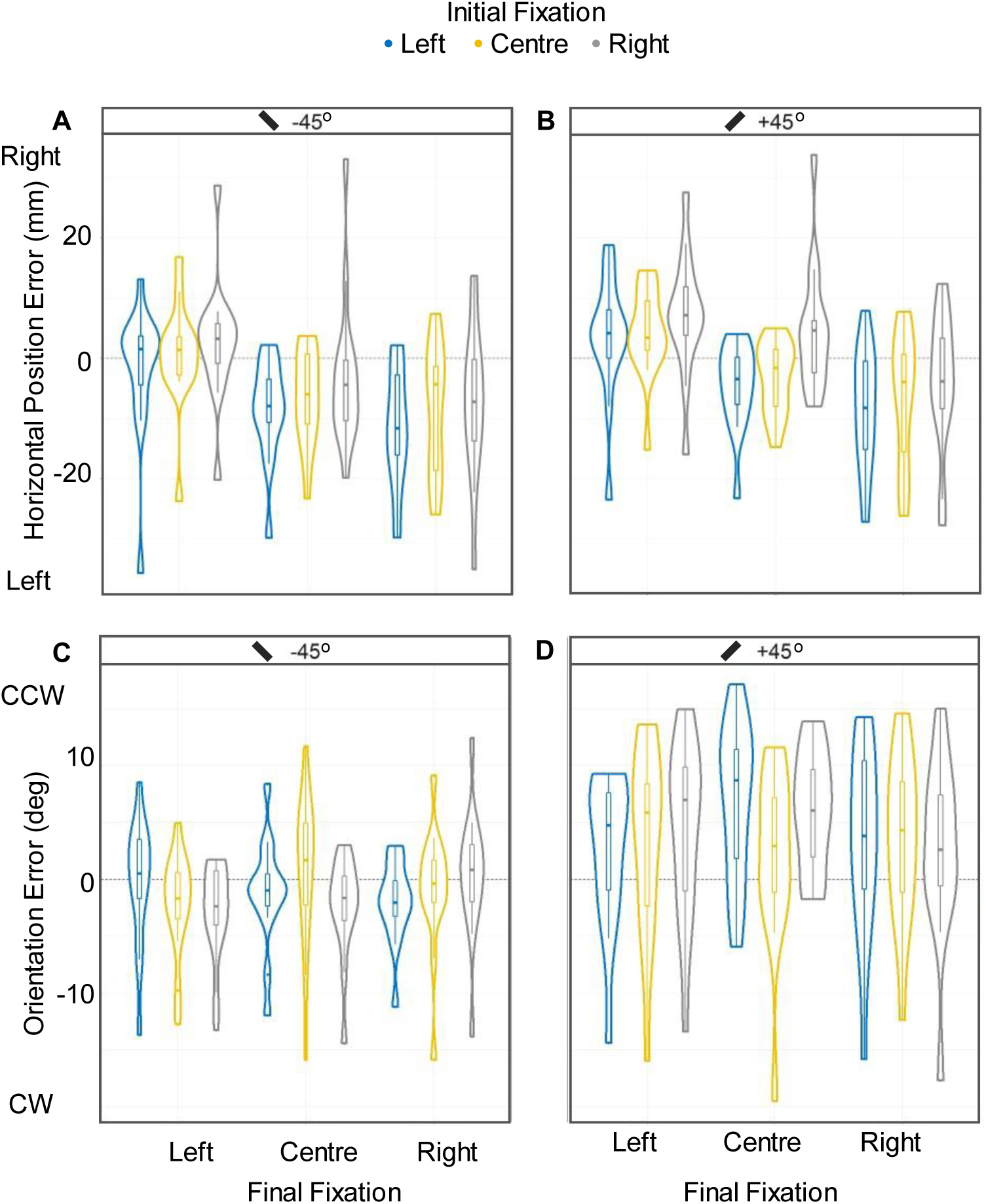
Distribution of Location and Orientation Errors for *Experiment 2*. Violin plots show the distri ution o t e artici ant’s average errors and include the median (depicted as the dot) and the interquartile range (as the bar in the center of the violin). Top panels: Violin plots representing placement location errors for (A) −45° target orientation and (B) +45° target template orientation. The data are grouped according to the final gaze locations and color-coded according to initial gaze locations. A positive value indicates movement towards the right, and a negative value indicates movement to the left. Bottom panels: Violin plots illustrating placement orientation for (C) −45° target template orientation and (D) +45° target template orientation. The values from the y-axis provide information regarding the direction of the error: a positive value indicates a counterclockwise rotation, and a negative value indicates a clockwise rotation. Other plotting conventions are the same as those for the upper row.

To investigate these trends, we conducted a repeated measure 3 × 3 × 2 ANOVA analysis, with three initial gaze locations (left, centre or right), three final gaze locations (left, centre or right) and two orientations (+45° or −45°) as the variables. A three-way ANOVA revealed a significant main effect main Final Gaze Location (F_2,26_ = 10.32, P < 0.001, η2 = 0.44 related to the gaze overshoot described above. There was also a significant main effect of Orientation on Placement Location (F_1,13_ = 20.16, P < 0.001, η^2^ = 0.61).

There was no significant main effect of Initial Gaze Location (F_2,26_ = 2.817, P > 0.5, η^2^ = 0.18), but Initial Gaze Location × Orientation showed a significant two-way interaction (F_2,26_ = 4.059, P < 0.05, η^2^ = 0.24). Post-hoc tests showed that (compared to saccades from centre gaze), saccades from leftward gaze (but not the rightward gaze) showed a stronger influence of both counterclockwise (mean difference = 4.55mm, d = 0.43) and clockwise (mean difference = 2.32mm, d = 0.20) orientation on placement location. There was no significant three-way interaction (Orientation × Initial Gaze Location × Final Gaze Location: F_4,52_ = 1.436, P > 0.05, η^2^ = 0.10).

These observations are further illustrated in Fig. 7 A, B and C, which show endpoint location errors as a function of final gaze, with initial gaze positions indicated by the symbols (left ▴, centre ●, rig t ▪ and template orientation indicated by the colour (orange rotated clockwise, blue rotated counterclockwise). In Fig. 7, the top row shows fixation data, the middle row centripetal / centrifugal saccade data, and the bottom row across saccade data. Fixation data show the typical ‘overshoot’ pattern, as do the centrifugal data and across data.

**Figure 7.**
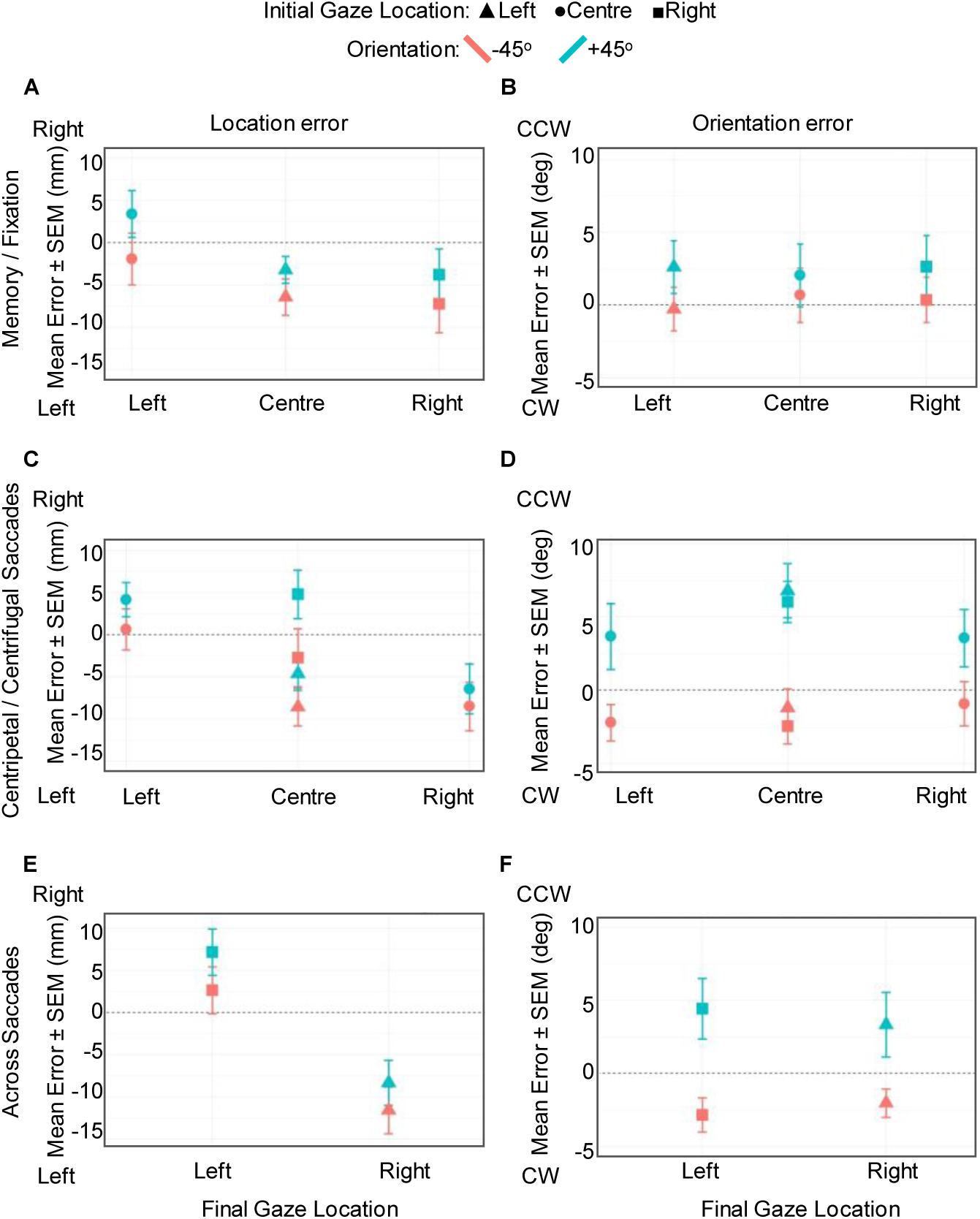
Mean Location Error (*left column*) and Orientation Errors (*right column*) in *Experiment 2*, plotted as a function of final gaze position and coded according to template orientation (−45° pink / +45° blue) and initial gaze location (left ▴, centre •, right ▪).*Top row*: *Fixation* data (where initial and final gaze location are identical); *Middle row*: *centripetal/centrifugal* saccade data; *bottom row*: *Saccade across* data.

The exception is the centripetal data, i.e., the saccades that end at centre in Fig. 7C. Here, placements with leftward saccades / clockwise templates and rightward saccades / counterclockwise templates hand small errors, as expected, but the opposite pairs (leftward / counterclockwise, rightward / clockwise) ended with large errors in opposite directions, as though overcorrecting for the placement overshoots observed at the original gaze position (in the fixation task). This accounts for the saccade / orientation interaction that emerged in our statistical analysis, and contradicts the normal rule that final gaze determines hand position errors.

#### Orientation error

A second ANOVA was conducted on the average difference between participants’ performed endpoint orientation and the actual target template orientation. Looking at the violin plot to see a summary statistic of the distribution of error, there is a greater variation and relative shift positive (counterclockwise) in the block final orientation for the +45° in comparison to the −45° (Fig. 6, C and D). A repeated measure 3 × 3 × 2 ANOVA analysis was conducted, with three initial gaze locations (left, centre or right), three final gaze locations (left, centre or right) and two orientations (+45° or −45°) as the variables. The three-way repeated measure ANOVA was conducted to investigate the influence of saccade types and target template orientation on endpoint orientation error.

The results revealed a significant main effect of template Orientation on hand orientation errors (F_1,13_ = 8.544, P < 0.05, ƞ^2^ = 0.4). There was no significant main effect of initial gaze (F_2,26_ = 0.488, P > 0.05, ƞ^2^ = 0.4) or final gaze location (F_2,26_ = 2.988, P > 0.05, ƞ^2^ = 0.19) on the endpoint orientation, but there was a significant two-way interaction between Initial Gaze Location and Orientation (F_2,26_ = 5.53, P < 0.01, η^2^ = 0.3) and a three-way interaction for Orientation, Initial and Final Gaze Location (F_4,52_ = 10.43, P < 0.000, η^2^ = 0.45). Follow-up tests showed no significant interaction between Initial Gaze Location, Final Gaze Location, and Orientation (all P > 0.05). In short, hand orientation errors were primarily influenced by template orientation, but this interacted with initial and final gaze location.

Fig. 7 B, D and F illustrate the influence of final gaze, initial gaze, and template orientation on object orientation using the same conventions as the left column, showing fixation data (top row), centripetal / centrifugal data (middle row) and across saccades (bottom row). A relative positive counterclockwise rotation was always seen for the +45° angle in the saccade conditions and a relative negative clockwise rotation for the −45° angle. In other words, people tended to overshoot the template rotation. However, this effect was quite small in the memory-fixation condition (Fig. 7B), compared to experiment one (with no delay) and compared to the even larger errors in the saccade data here (Fig. 7, D and F). These subtle differences observed in the patterns likely correspond to the interactions mentioned earlier, but there is no clear main effect of gaze endpoint on orientation, i.e., the data do not consistently slope to the left or right.

## DISCUSSION

Whereas the influence of vision and gaze position on reach / pointing location is well studied, their influence on object placement (in both the location and orientation domains) has largely been ignored. In this study, we investigated the influence of gaze location, level of visual feedback, saccades, and target template orientation on the accuracy of orientation and horizontal position in an object placement task. In Experiment 1, we assessed the accuracy in fixation gaze conditions, with and without visual feedback of the target template and hand location during the placing movement. As in reaching and pointing movements (2–4), there was a systematic overshoot of endpoint location in the direction opposite to the gaze location that increased in the open-loop (no visual feedback) condition. Participants also tended to overestimate the rotation of the placement template, and this pattern was influenced by gaze location and visual feedback. We also tested (Experiment 2) how an increased memory delay and various saccade patterns would affect placement accuracy. While a consistent effect of gaze location could be observed in endpoint location during placement, it was not observed in orientation for all conditions. However, endpoint orientation was more influenced by the target template orientation than the gaze location.

### General Bias in Placement Location

Our results showed a general bias in placement endpoint location, most easily observed when participants looked straight ahead at the horizontal template. It is possible that this was a visual strategy to allow participants to see the edge of the template as the block approached its final destination. Alternatively, this could be linked to a proprioceptive motor bias. Similar proprioceptive bias in the endpoint reach has been shown to have a gaze-independent component (41). In either case, one might expect this bias to reverse for the left hand (this was not tested).

### Influence of Template Orientation on Placement Orientation

A consistent observation in both experiments is that participants tended to overestimate rotations of the stimulus template from the horizontal. Since we only used template orientations that participants reported to be comfortable with, and the effect was an overshoot rather than undershoot, these effects were unlikely to be due to biomechanical constraints. A similar effect of target template orientation in the endpoint rotation error was reported in grasp such that the grasp angle depended on the object orientation (42, 43). In addition, Mason & Bryden (44) reported a similar over-rotation in a grasp, which was more pronounced for the oblique orientation. It is possible that this was a deliberate visual strategy (i.e., to avoid obscuring the template). However, this does not explain why these patterns were nearly identical in both the open-loop and closed-loop conditions. This suggests either a learned strategy or a miscalibration in the ‘dorsal stream’ transformations for action.

### Influence of Gaze Position on Placement Location

Reaching and pointing studies have shown a gaze-dependent overshoot effect in the direction opposite to the gaze location before movement onset (3, 18, 19, 45, 46). These original findings have been extended across different memory delays, with egocentric and allocentric frames of reference (2, 4, 47–49). Finally, such gaze-dependent errors interact with the errors related to initial hand position and may be magnified by damage to the parietal cortex (23, 24).

Regarding object placement, we observed similar gaze-centered patterns of reach error during placement. This suggests that the overshoot effect presents various types of goal-directed movement and is not exclusive to a simple reach movement. Presumably, these errors result from miscalibrations in the gaze-dependent transformation from visual coordinates to the motor coordinates for hand motion (50, 51). Two specific explanations for this phenomenon are that the target direction is overestimated relative to gaze (3) or that gaze direction is underestimated relative to the target (52). An alternative theory, supported by some evidence, is that the error does not occur in the reach transformation, but rather in the transformation of hand position signals into gaze-centered coordinates, when compared to vision for computation of the reach plan (53–56). In any case, the observation that this effect generalizes to different behaviors (pointing, reach, placement) suggests a common computational mechanism.

### Influence of Gaze Position on Placement Orientation

In the current study, we observed gaze-dependent orienting errors that changed direction based on the target template location and orientation. This suggests that gaze location and target template orientation are two elements that interact and influence the final orienting error observed in object placement. As in grasp, the gaze-dependent orienting error in placement was more pronounced for oblique than cardinal orientations (32, 33). This might reflect a tendency to process reach information along coordinate system-like channels (57), although little physiological evidence supports this idea (58).

It has been shown that gaze direction interacts with 2D wrist pointing direction for grasp orientation (27), but it is less clear why gaze direction would interact with 3D orientation of the hand about the axis of the wrist (as in the current task). Object location and orientation are spatially independent variables that do not tend to cross-correlate in visuomotor adaptation studies (30). Further, orienting objects in this dimension involves coordinated movements of the fingers, wrist, and upper arm (28). These factors again suggest a high-level but non-specific gaze-orientation interaction, perhaps in the dorsal visual stream. We will speculate about such mechanisms in a later section below.

### Influence of Visual Feedback and Memory Delay

Previous studies have shown that having visual feedback of the target and/or hand position during reach influences accuracy in the movement, presumably by aiding the online calculation of the reach vector (55, 56, 59). Conversely, prolonging the delay period following a transient visual stimulus beyond 250ms decreases movement accuracy, presumably due to degradation in the egocentric visual memory of the target stimulus (36, 60, 61).

To determine if these factors held during placement movement, we investigated how changes in the presence of visual feedback influence participants’ accuracy in their final placement of an object. We first looked at the influence of visual feedback on gaze-dependent errors in Experiment 1. Consistent with previous studies (2, 3, 62), we found a significant increase in gaze-dependent location overshoots in the absence of visual feedback. In contrast, visual feedback did not influence gaze-dependent orientation errors. This suggests that these orientation errors were not due to a visual strategy but rather some internal visuomotor miscalibration, perhaps related to gaze-dependent transformations in the ‘dorsal stream’.

As expected, gaze-dependent location errors persisted with the addition of a memory delay in Experiment 2. This remains consistent with the notion of dorsal stream errors that appear in the absence of visual feedback, which then requires a shift to ventral stream processing (63). But surprisingly, gaze-dependent orientation errors were dramatically reduced during memory / fixation condition (Fig. 7B). It has been suggested that the addition of a memory delay reach tasks engages ‘ventral stream’ visual processes (64, 65). If the ventral stream is better at processing orientation information than the dorsal stream, this might explain why orientation errors were reduced in this condition. This functional anatomy is highly speculative, but our behavioral results highlight that memory-guided reach may rely on different neural mechanisms and this does not always lead to inferior performance.

### Influence of Saccades

Previous studies have shown that in the presence of saccades, the final gaze position determines memory-guided reach errors, and this has been used to support the notion of gaze-centered updating (3, 4, 51, 66). However, those studies consistently relied on centrifugal saccades (from the target toward some peripheral fixation position). In the real world, saccades may follow a variety of different repertories. The present study examined three of these repertoires (centrifugal, centripetal, across target template) to see if memory-guided placement shows the same pattern (i.e., dependence only on final gaze), and if those results generalized to different saccade amplitudes, directions, and positions.

Our overall statistical analysis confirmed that final gaze position, rather than initial gaze position, determined the influence of saccades on memory-guided placement. However, there were more nuances when we examined specific types of saccades. Centrifugal saccades and saccades that crossed the target template followed the expected pattern, supporting the notion that gaze-centered updating for memory-guided reaching also generalizes to placement tasks. The main difference was in centripetal saccades, which appeared to ‘overcorrect’ the influence of initial gaze position, i.e., produce overshoots opposite to the initial gaze position, depending on the saccade direction and the orientation of the template. Further, the presence of a saccade in the memory interval re-introduced the orientation overshoots observed in experiment one, perhaps (if the ventral stream memory theory is correct) due to some re-setting of dorsal stream representations during updating (22, 23, 64).

### Caveats and limitations

As in any such experiments, the results in this study are specific to the task and can only be generalized to other tasks with further experimentation. Clearly, this was a visually impoverished environment with artificially fixed gaze positions; thus, the errors reported here might be suppressed during gaze fixation in a richer environment containing allocentric cues (36, 67). However, gaze is not always locked to placement targets in real-world circumstances (68), and peripheral vision does not have the same acuity. Another caveat is that, although our participants interacted with a real 3D object, they placed it toward a 2D surface that did not have depth or somatosensory cues. In our increasingly technology-dominated lives, people often interact with 2D touch screens, but it is possible that additional gaze-dependencies and brain mechanisms are engaged in a complete 3D environment (69–74). Finally, although this study was inspired by the literature on eye-hand coordination during reaching / pointing, we did not directly compare reach and placement in the current experiment, so could only provide qualitative comparisons based on the literature.

### Possible Neural substrates

The literature on neural control of goal-directed action is dominated by reach and grasp experiments, whereas relatively few studies have examined brain activity beyond goal acquisition. Several functional Magnetic Resonance Imaging studies have investigated the neural mechanisms of object manipulation. For example, the size-weight illusion has been used to investigate the neural correlates of force correction and weight prediction (75). Object manipulations such as these involve the recruitment of regions in the posterior parietal cortex (PPC), the cerebrum, and the motor cortex.

Thus, at this time, one must speculate on the neural mechanisms of placement based on what is known about the reach / grasp system. As noted above, the dorsal visual stream, corresponding dorsal occipital and posterior parietal cortex is the likely site for the integration of visual and other sensory information to guide visuomotor plan (31, 76, 77). Mid-posterior intraparietal cortex (mIPS) is involved in calculating the hand-target vector for reach transport (78, 79), and thus might likewise be engaged in our match-to-template task. Anterior IPS are involved in grasp formation (76), and thus might be involved in the ‘inverse grasp’ for placement: letting go of the object at the end. The superior parietal occipital cortex (SPOC) has been shown to have both transport and grasp signals (e.g., 80, 81), and is specifically involved in orienting the hand (82); so again, it might be engaged in a task like this. Of course, PPC does not operate in isolation but rather communicates extensively with frontal reach areas such as the dorsal and ventral premotor cortex and primary motor cortex (31, 79, 83).

Although the dorsal stream is associated with the use of online visual control, lateral and superior occipital cortex and PPC have been implicated in memory and planning for reaching in a delay task (56, 84, 85). These areas are also implicated in the gaze-dependent transformations for reach (58, 86–88). However, as noted above, it has been argued that the ventral stream becomes engaged in memory-guided reaches, especially as the delay increases (63, 89). Our observations show that gaze-centered updating is involved in placement, a phenomenon associated with PPC (22, 76). Specifically, we recently implicated a cortical network involving frontal eye fields, inferior parietal cortex (specifically supramarginal gyrus), aIPS, and dorsal PPC in updating grasp orientation during saccades, which may have played a similar role here (35).

## Conclusions

Prehension and placement are both goal-directed movements that require identifying the target location and its orientation in order to successfully perform the desired movement. However, they differ greatly in the intention, as one (reach) is generally used to acquire an object, and the other (placement) is used to dispose of it. It does, however, appear that they follow certain similar kinematics rules of eye-hand coordination. As in reach movements, placement movements (in our task) showed degraded performance in the absence of vision and a trend to overestimate direction relative to gaze. The influence of gaze on orientation was less systematic, but we observed orientation overshoots in our task as well as various subtle interactions between gaze and orientation, depending on different levels of visual feedback and memory delay.

These observations suggest that at least some aspects of the acquisitional reach system (such as early gaze-centred processing) are shared by the placement system and provide other details that need to be accounted for in modeling the system. They also have practical implications. If our observations generalize to other conditions, this would have consequences for any technology or task that requires highly accurate placement in both the location and orientation on a two-dimensional plane. Further, this study provides a baseline of health behavior for further studies in patients. For example, based on previous reach and pointing studies, one might expect these errors to increase in any pathology that impairs parietal cortex function (23, 24).

## ACKNOWLEDGMENTS

The authors thank Saihong Sun for technical support.

## AUTHOR CONTRIBUTIONS

G.N.L, J.D.C, X.Y. and E.F. conceived and designed research; G.N.L. performed experiments; G.N.L analyzed data; G.N.L., J.D.C and E.F. interpreted results of experiments; G.N.L. prepared figures; G.N.L. and J.D.C. drafted manuscript; G.N.L., J.D.C., E.F. and X.Y. edited and revised manuscript; G.N.L., J.D.C., E.F. and X.Y. approved the final version of manuscript.

